# Acute memory deficits in chemotherapy-treated adults

**DOI:** 10.1101/215731

**Authors:** Oana C. Lindner, Andrew Mayes, Martin G. McCabe, Deborah Talmi

**Affiliations:** Division of Neuroscience and Experimental Psychology, School of Biological Sciences, University of Manchester, Zochonis building, Oxford Road, Manchester, M139PL.; Division of Molecular & Clinical Cancer Sciences, Faculty of Biology, Medicine and Health, University of Manchester

**Keywords:** memory, forgetting, cognition, cancer, chemotherapy

## Abstract

Data from research on amnesia and epilepsy are equivocal with regards to the dissociation, shown in animal models, between rapid and slow long-term memory consolidation. Cancer treatments have lasting disruptive effects on memory and on brain structures associated with memory, but their acute effects on synaptic consolidation are unknown. We investigated the hypothesis that cancer treatment selectively impairs slow synaptic consolidation. Cancer patients and their matched controls were administered a novel list-learning task modelled on the Rey Auditory-Verbal Learning Test. Learning, forgetting, and retrieval were tested before, and one day after patients’ first chemotherapy treatment. Due to difficulties recruiting cancer patients at that sensitive time, we were only able to study 10 patients and their matched controls. Patients exhibited treatment-dependent accelerated forgetting over 24 hours compared to their own pre-treatment performance and to the performance of control participants, in agreement with our hypothesis. The number of intrusions increased after treatment, suggesting retrieval deficits. Future research with larger samples should adapt our methods to distinguish between consolidation and retrieval causes for treatment-dependent accelerated forgetting. The presence of significant accelerated forgetting in our small sample is indicative of a potentially large acute effect of chemotherapy treatment on forgetting, with potentially clinically-relevant implications.

## Introduction

Many believe that early long-term memory (LTM) is initiated by rapid consolidation triggered at encoding and that it lasts between ten minutes and several hours (Kandel, Dudai, & Mayford, 2014; McGaugh, 2000). More slowly-triggered consolidation then enables memories to last for hours, days, or longer (Wixted, 2004). Research in animal models has revealed double dissociations between rapid and slow consolidation and provided substantial evidence that drug-induced de-novo protein synthesis disruption impairs slow consolidation and accelerates forgetting over 24 hours (Kandel et al., 2014; McGaugh, 2000). By contrast, there is only sparse evidence for dissociation between fast and slow consolidation in humans. Here we report results from the first study of the cognitive effects of acute cancer chemotherapy treatment in humans, providing preliminary for this dissociation.

The canonical human memory literature does not currently distinguish between minutes-long‘early’ and hours-or-days long ‘delayed’ LTM. This is partly because human forgetting curves (which depict performance on LTM tests as a function of time since encoding) are typically monotonically decreasing, and described well by a power function (Kahana & Adler, 2002; Rubin et al., 1996; Wixted, 1990). The single dissociation between early and delayed LTM is well established. There is ample evidence that the hippocampus is key to memory persistence, although exactly what that entails is still actively debated (Carlesimo, Cherubini, Caltagirone, & Spalletta, 2010; Dewar, Della Sala, Beschin, & Cowan, 2010; Gershman, Blei, & Niv, 2010; Hardt, Nader, & Nadel, 2013; Kopelman et al., 2007; Mayes & Roberts, 2001). Damage to the hippocampus, other medial temporal lobe structures and their connections, leads to an accelerated loss of the ability to freely recall recent inputs. Free recall performance is decreased when tested within the first ten minutes, after which free recall is usually poor (Isaac & Mayes, 1999a, 1999b). Evidence for the reverse dissociation is less well-established. Some patients with temporal lobe epilepsy show normal free recall within the first ten minutes after encoding, but accelerated long-term forgetting (ALF) as indicated by free-recall measures in delayed tests (Alber, Della Sala, & Dewar, 2014; Elliott, Isaac, & Muhlert, 2014; Hoefeijzers, Dewar, Della Sala, Butler, & Zeman, 2015; Isaac & Mayes, 1999a, 1999b). The strength of these dissociations between organic amnesia and ALF is disputed, but if replicated, these findings support a distinction between human rapid and slow LTM consolidation.

Determining whether there is more than one kind of consolidation in humans might be helped by directly manipulating the putative neurobiological processes, using distinct pharmacological treatments. Clearly, we cannot administer drugs to humans that may cause amnesia. In this study, we took a novel approach to address this challenge, by examining, for the first time, rapid and slow consolidation in non-Central Nervous System (nCNS) cancer patients who have undergone chemotherapy.

Cancer chemotherapy treatments are interesting for the study of memory consolidation for two reasons. First, chemotherapy treatment involves the delivery of drugs that are very toxic to humans. While not all cytotoxic drugs cross the blood-brain barrier, pro-inflammatory cytokines that are triggered by the treatment can reduce the protective capabilities of the barrier, allowing some of the cytotoxic drugs to cross it (Coussens & Werb, 2002; Pan et al., 2011; Terrando et al., 2010). The CNS effects of cytotoxic drugs may be due to the disruption of the blood-brain barrier itself, increase of cytokine expression in the CNS, or oxidative stress (Ahles & Saykin, 2007; Seigers & Fardell, 2011). For instance, while anthracyclines do not readily cross the blood-brain barrier (da Ros et al., 2015), in animals they have nevertheless been linked to apoptosis, inhibition of neuro-and gliogenesis (Dietrich, Prust, & Kaiser, 2015; Kaiser, Bledowski, & Dietrich, 2014), and reductions in serotonin-induced long-term synaptic facilitation, required for the activation of LTM consolidation processes (Liu, Zhang, Coughlin, Cleary, & Byrne, 2014). These neurobiological effects have consequences for memory; for example, suppression of neurogenesis in the medial temporal lobe disrupts hippocampus-dependent memories (Arruda-Carvalho, Sakaguchi, Akers, Josselyn, & Frankland, 2011; Luu et al., 2012; Sahay et al., 2011; Zhang, Zou, He, Gage, & Evans, 2008). Indeed, research in animals shows that even the first chemotherapy treatment impairs memory; impairments can increase in severity as treatment progresses (Rzeski et al., 2004) and are associated with neuronal damage (Dietrich et al., 2015; Kaiser et al., 2014).

Second, although the acute effects of cancer chemotherapy treatment on memory are not known, it is well established that cancer survivors treated with chemotherapy exhibit chronic structural and functional brain damage and cognitive difficulties. This literature is heterogeneous, but most cross-sectional behavioural studies, recent longitudinal studies (Janelsins, Kesler, Ahles, & Morrow, 2014), and structural and functional imaging evidence (de Ruiter & Schagen, 2013; Deprez, Billiet, Sunaert, & Leemans, 2013; Pomykala, de Ruiter, Deprez, McDonald, & Silverman, 2013) have consistently demonstrated chronicimpairments in memory,executive functions, and the brain regions that subserve them. Declarative memory is one of the cognitive functions most frequently affected (Lindner et al., 2014; Saykin, de Ruiter, McDonald, Deprez, & Silverman, 2013). Crucially, these behavioural markers and structural/functional brain changes are independent of anxiety, depression, or any other patient-reported outcomes.

Taken together, cancer chemotherapy treatment is thought to induce changes in the brain that impair memory acutely and chronically. This presents an interesting opportunity to study consolidation causally, under controlled conditions, in a population without neurological or psychiatric history. Beyond this theoretical interest, evidence for the acute cognitive effects of chemotherapy has obvious clinical relevance, with potential impact on patient-doctor communication practices. The reason the effects of acute chemotherapy on cognition have not been studied until now has to do with the obvious challenges of conducting experiments during a very sensitive time for patients, within weeks after diagnosis and around the onset of a difficult treatment. We report the results of a small study, with 10 non-CNS cancer patients (and their matched controls) where we examined 2-minute and 24-hour delayed memory before, and immediately after patients’ very first chemotherapy treatment. We hypothesised that patients will exhibit accelerated forgetting following their first treatment, both relative to their pre-treatment performance and to that of matched controls. We suspected that cancer treatment may disrupt de-novo protein synthesis and hence impede slower memory consolidation whether or not learning and/or retrieval are also disrupted. Our results suggest that slow consolidation is impaired in this group, in agreement with our hypothesis, but the small sample size means that they can only be considered preliminary. We report them in order to encourage larger studies of this topic.

## Methods

### Participants

The study was approved by the National Research Ethics Services Committee North West. Exclusion criteria common to all participants included a previous history of cancer and/or chemotherapy, hormonal treatment, cranial irradiation, brain injury, a history of mental health problems or substance abuse, previously exposed to mood altering drugs, or if they were not proficient in English.

#### Patients

Cancer patients were recruited to the study between November 2011 and April 2014. Patients were approached by their clinical team if they were between 16 and 50 years old and had been diagnosed with one of four cancers most prevalent in working ageyoung and middle aged adults (CRUK, 2014): sarcoma, lymphoma, breast cancer, or germ cell tumour. Because the mechanism with which cytotoxic drugs disrupt the CNS are varied and ultimately not yet known, and because most chemotherapy regimens involve the administration of multiple cytotoxic agents, we did not limit inclusion to patients receiving drugs known to cross the blood brain barrier. To be included in the study, patients had to be available to be evaluated before their adjuvant or neoadjuvant treatment.

Figure 1 depicts the recruitment process. The self-exclusion of unwell patients suggests that the participants who completed Session 3 may have had a better health and emotional status compared to decliners. Due to logistical difficulties, three participants could not be tested using a computer in either Sessions 2 or 3. To be cautious, the final sample only includes patients who were tested on the computerised version of the task, but we also comment on any differences between the analyses of the two samples. All patients in the final sample had received antiemetics as part of their first treatment. None had additional medical co-morbidities. All women were pre-menopausal. There were no differences in the distribution of diagnoses, demographic details, neuropsychological characteristics, or patient-reported outcomes between the patients who took part in this study and those who did not (Supplementary Tables S1-S2).

**Figure 1.**
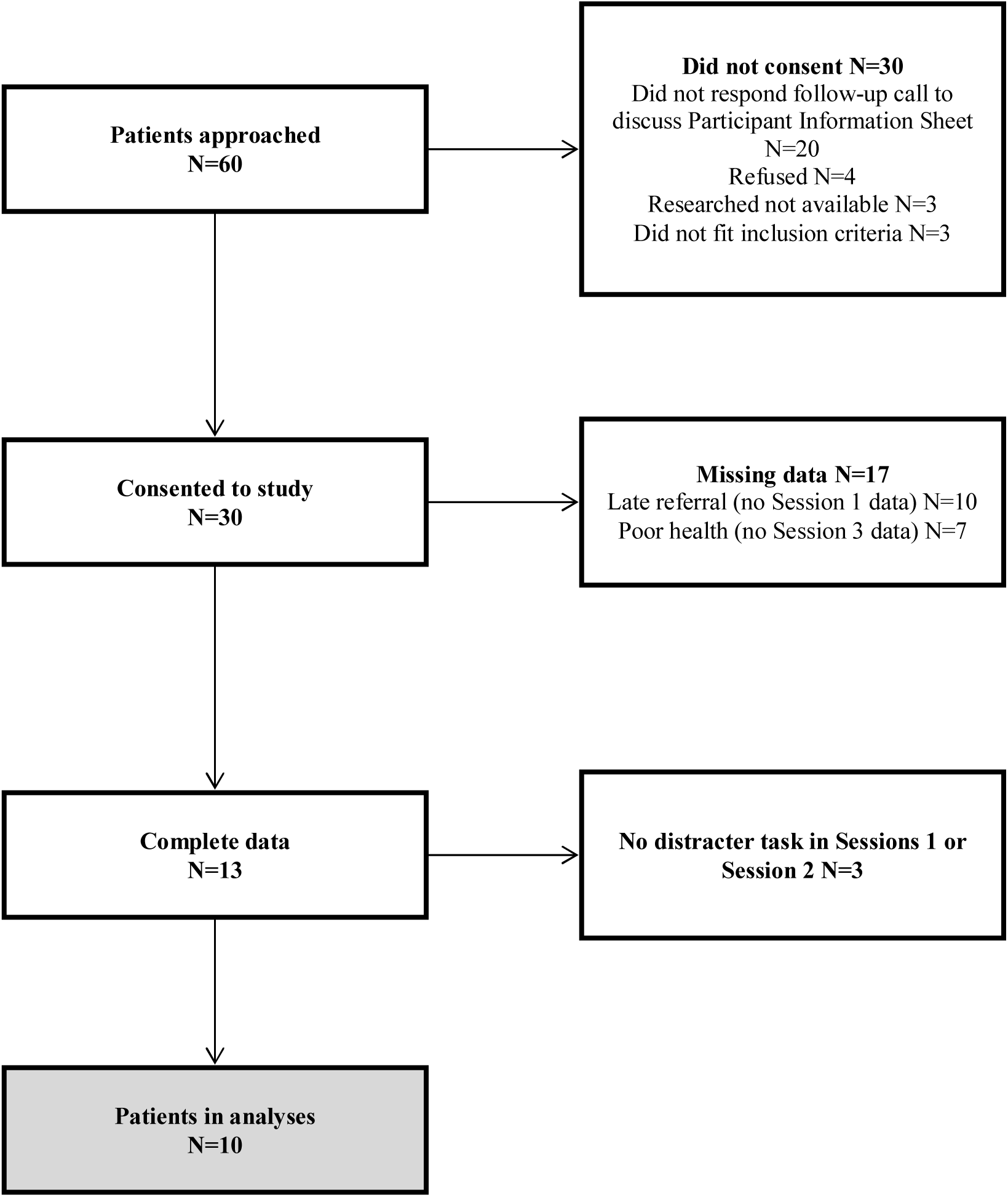
Flowchart of patients included in study and analyses.

#### Controls

Control participants (N=10), recruited through adverts, were matched to patients on education, sex, and age (+/− 5 years). There were no significant differences between participants on any demographic variables (Table 1).

**Table 1.** Sample Characteristics

| Patients |  |  |  |  |  | Controls |  |  |
| --- | --- | --- | --- | --- | --- | --- | --- | --- |
| ID | Diagnosis | Treatment(Cycles) | Age | Sex | Education | Age | Sex | Education |
| 1 | Ewing sarcoma* | VIDE (6)<br>VAI (8) | 20 | M | College | 24 | M | Degree |
| 2 | Osteosarcoma | MAP (4) | 20 | F | College | 19 | F | College |
| 3 | Germ cell tumour | BEP (3) | 30 | M | Degree | 30 | M | Degree |
| 4 | Osteosarcoma | MAP (6) | 17 | M | College | 20 | M | Degree |
| 5 | Hodgkin lymphoma | ABVD (2)<br>AVD (4) | 19 | F | College | 20 | F | College |
| 6 | Ewing sarcoma* | VIDE (6)<br>VAI (8) | 17 | F | College | 21 | F | College |
| 7 | Breast cancer | FEC-T (6) | 46 | F | Degree | 46 | F | College |
| 8 | Breast cancer | FEC-T (6) | 45 | F | Degree | 46 | F | Degree |
| 9 | Breast cancer | FEC-T (6) | 45 | F | Degree | 46 | F | Degree |
| 10 | Hodgkin lymphoma | ABVD (6) | 22 | F | Degree | 20 | F | College |
| M | 30% Breast cancer |  | 28.1 | 70% F | 50% | 29.2 | 70% F | 50% |
| (S | 40% Sarcoma |  | (12.44 | 30% M | College | 0 | 30% | College |
| D) | 20% Hodgkin's lymphoma |  | ) |  | 50% | (12. | M | 50% |
| % | 10% Germ cell tumour |  |  |  | Degree | 01) |  | Degree |

### Instruments

#### Word lists

Five different word lists were created in a pilot study using 60 young adults. The words consisted of concrete nouns from the Snodgrass and Vanderwart (1980) database. To limit proactive interference and list confusions each list contained 24 words from two categories representing one natural and one man-made concept. Category names were used as semantic cues for free recall in the beginning of each Session. Lists were equivalent in familiarity and word frequency and were 4-10 letters in length (Supplementary Table S3). The first two letters of each word were unique and served as a cue in the cued recall test. Three lists were selected for each patient and one list allocated to each Session through a balanced Latin square method (Reese, 1997).

#### Distracter task

The task consisted of two similar pictures containing 15 differences. Participants were asked to find as many differences as they could within 2 minutes.

#### Additional tests

Education is often used as a proxy of general intellectual performance to enable matching between groups (Neisser et al., 1996), although this may not be sufficient in between-group cognitive studies (Deary & Johnson, 2010). Following on from the discussion in Lindner et al. (2014) we attempted to match the groups better by measuring full-scale IQ through the Wechsler Test of Adult Reading (WTAR, Strauss et al., 2006), a good estimate general cognitive functioning. To control for potential confounders and adhere to the recommendations of the International Cognition and Cancer Task Force (ICCTF, Wefel et al., 2011) we attempted to evaluate patients’ neuropsychological status with standardised measures. We also asked participants to fill in several questionnaires at home and send them back to us in a self-addressed envelope. Logistic difficulties, inherent to clinical settings, meant that not all patients were able to complete all the neuropsychological and patient-reported outcomes measures. Available results (Supplementary Table S2) suggest that our sample generally exhibited low level of emotional distress, and were not very different from controls on learning and memory measures. All participants completed the WTAR, which we were therefore able to include as a covariate in the analyses.

### Procedure

A difficulty in carrying out repeated memory tests in clinical settings is that instruments recommended in neuropsychological studies of cancer patients (Vardy, Wefel, Ahles, Tannock, & Schagen, 2008) cannot longitudinally assess forgetting across a 24-hour period, a feature we particularly wanted to examine. To address this, we designed a new word-learning task. The specific testing procedure, has a strong theoretical justification to enable the investigation of learning, forgetting, and retrieval processes. Figure 2 provides a graphical summary of the procedure. The task was administered in three Sessions, one per day,over three consecutive days. Patients received the first treatment after the first two Sessions and before the third. Sessions were short (10-15 minutes) to facilitate the administration of this non-routine test before and after patients’ first treatment.

**Figure 2.**
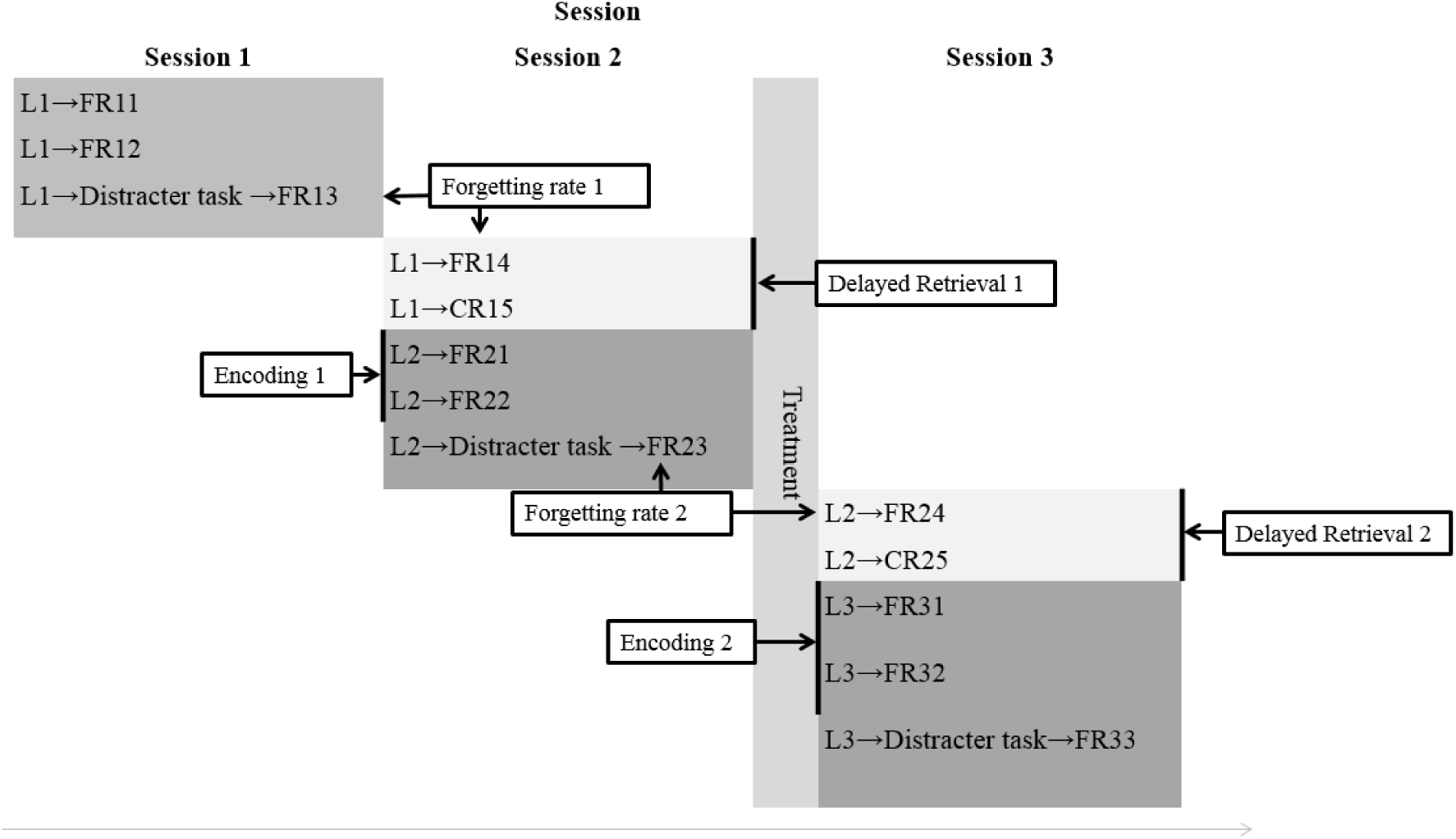
Memory task procedure. Each Session corresponds to a day of testing, on three consecutive days of testing. In patients, Sessions 1 and 2 take place before treatment and Session 3 after treatment. L: On-screen presentation (2.5 seconds) of each word in Lists 1, 2, 3 (participants had to include each word in a verbalised sentence). FR11, 12, 13, 14: three immediate and a delayed oral Free Recall Test of List 1. FR21, 22, 23, 24: three immediate Free Recall Tests and a delayed free recall Test of List 2. FR31, 32, 33, 34: three immediate Free Recall Tests of List 3. In all Sessions the third free recall takes place after a 2-minute distracter task. CR15, CR25: Cued recall tests of Lists 1 and 2. Forgetting rate 1 and 2: proportion of information forgotten between 2-minute and 24-hour delayed FR tests before (FR13 and FR14) and after (FR23 and FR24) treatment. Encoding 1 and 2: learning performance between the two immediate FR tests before (FR21 and FR22) and after (FR31 and FR32) treatment. Delayed Retrieval 1 and 2: proportion of information recalled between the 24-hour delayed free and cued recall tests before (FR14 and CR15) and after treatment (FR24 and FR25).

The task was modelled after the Rey Auditory Verbal Learning Test (RAVLT, Strauss, Sherman, Spreen, & Spreen, 2006). It was short and flexible enough to be administered to patients before and immediately after their first treatment, and sensitive enough to capture and differentiate between potentially mild learning, consolidation, and retrieval deficits, which could all underlie impaired memory (Mayes, 1995; Mayes & Roberts, 2001). The differences between our task and RAVLT were the use of categorised words (limiting interference in a delayed recall context); the increased number of items and reduced number of learn/recall trials (limiting ceiling effects); the inclusion of an unrelated distracter task before the final free recall (limiting recall from working memory and potential interference inherent to verbal tasks); and the use of a cued recall test to tap into memory performance under aided retrieval conditions (avoiding potential ceiling effects expected in a recognition test). The exact instructions used for each session are provided in the Supplementary Material.

In each Session, each word list was studied, tested through free recall, studied again, and tested again through free recall. During study, each word was presented on a screen for 2.5 seconds. Participants produced a sentence out loud with the target word (e.g. “The **helicopter** is in the sky”), after which they pressed a key to proceed to the next word. Sentences were not recorded but participants adhered to the specific task instruction of not using the same sentence for different words; sentences could be the same for each learning occasion. Recall trials were terminated if participants stopped verbally recalling items for more than 20 seconds. The experimenter recorded both the words and intrusions produced by participants. After the second free-recall test, the list was studied a third time, the distractor task was administered, and the list was tested again. Memory for each list was tested a fourth time in the next Session, 24 hours later. Finally, each list was tested for the fifth time with cued recall.

#### Testing flexibility

Evaluations were performed non-routinely and at a sensitive time for patients hence they had to be flexible, while maintaining an appropriate experimental control. Patients and controls completed the task either in the hospital (university, respectively) or at home, while using the same procedures (i.e. if a patient was tested at home in Session 2 they were matched to a control who was also tested at home). When at home, participants were tested using a CD on their own computer, while speaking to the experimenter over the telephone. They entered a code to access the program, ensuring participants were not exposed to the material prior to testing.

### Analyses

An accelerated forgetting rate may be present on its own, or together with learning and/or retrieval deficits. As the focus of this study is on forgetting, we will discuss that measure first. Raw task performance scores are provided in supplementary Table S4.

#### Forgetting

Consolidation disruptions could be measured as an increase in forgetting rates (Averell & Heathcote, 2011; Wixted, 2004). It is difficult to interpret the raw number of words forgottenwithout accounting for how many words were remembered initially (Loftus et al., 1985). Consider two participants who recall 10 and 15 words in the early test, and 5 and 10 in the late test. Both forgot 5 words, representing 50% and 33% of the words initially recalled by the first and second participant, respectively. Forgetting rates were computed by dividing the number of words recalled in the later test by the number of words recalled in the earlier test, multiplied by 100. This measure compensates for initial recall levels and their potential drivers, including contextual emotional distress, learning problems, or difficulties with working memory, attention and concentration, factors which are therefore unlikely to affect forgetting rates.

Changes in rapid consolidation were explored by measuring *immediate forgetting rates*, namely, the difference between the second and third recall tests before (Session 2, (FR23-FR22)/FR22*100) and after treatment (Session 3, (FR33-FR32)/FR32*100). Changes in slow consolidation were explored by measuring *delayed forgetting rates*, namely the difference between the third and fourth recall tests, which were separated by 24 hours: once when both the study and the test occurred before treatment (Session 2 vs. Session 1, (FR14-FR13)/FR13*100) and once when study occurred before treatment, but test occurred after treatment (Session 3 vs. Session 2, (FR24-FR23)/FR23*100).

#### Learning

Learning scores were computed by averaging the percentage of words recalled in the two immediate free-recall tests of the same list. Two learning scores were computed: before treatment (Session 2, averaging FR21 and FR22) and after treatment (Session 3, averaging FR31 and FR32).

#### Retrieval

Retrieval indices included retrieval scores and intrusions. Retrieval deficit were suggested if cued recall improved performance relative to free recall more in patients than in controls, because cued recall reduces the need for organised search during retrieval by providing an aid. Scores were computed as the proportion of stimuli retrieved in the delayed cued recall relative to the preceding delayed free recall test, multiplied by 100. Two retrieval scores were computed: before treatment (Session 2, (CR15-FR14)/FR14*100) and after treatment (Session 3, (CR25-FR24)/FR24*100). Similarly to forgetting rates, the retrieval scores cannot be affected by initial learning difficulties, although lower retrieval scores could indicate problems with executive control. Excessive intrusions could suggest non-adherence to task instructions and a potential frontal-dependent memory disruption (Baddeley & Wilson, 1988).

#### Statistical analysis

Learning, immediate and delayed forgetting rates, and delayed retrieval rateswere normally distributed, as evaluated with the Shapiro-Wilk test (Shapiro, Wilk, & Chen, 1968). Data was analysed with repeated ANOVAs with the factors Group (patients/controls) and Session (before/after treatment). For brevity, only significant results are reported (p<.05). We examined forgetting with two planned one-tailed t-tests of the difference between patients and controls following treatment, and for the difference between patients before and after treatment. For all normally distributed data we report Hedge’s g effect size and its corresponding 95% confidence interval (CI, Borenstein, Hedges, Higgins, & Rothstein, 2009). Intrusions produced before and after treatment were not normally distributed, and were analysed with Mann-Whitney tests.

#### Statistical Power

Recruitment difficulties reduced our planned sample substantially. We hence ran a compromise power analysis, in which the power is evaluated as a function of effect size (set at minimum .20), sample size (N=10 per group), and an error-probability ratio (β/α) of 1 (i.e. equal probability of obtaining a difference through error or otherwise). It yielded a 71% probability of detecting small differences in our sample and a 61% probability to detect small differences if one covariate were included in the analysis (Faul, Erdfelder, Lang, & Buchner, 2007).

## Results

### Group characteristics

The three sessions took place approximately 24 hours apart. There were no differences between groups on session-to-session intervals (first delay, t_18_=1.95, p>.05; second delay, t_18_=−.12, p>.05). On the distracter task, no participant in either group identified all the differences in the allocated time.

Matching groups on education resulted in an equal number of controls and patients with college or university degrees. Despite that, patients had a lower FSIQ than controls, albeit still in the normal range (t_18_=−3.02, p<.01). Because these scores have a moderate relationship with the Memory Quotient of the Wechsler Memory Scale, the lower score in patients could give rise to impairment in our task. By controlling for FSIQ, we account for any potentiala-priori performance disadvantages in patients relative to controls. Similarly, covarying FSIQ could remove important variance because of these expected correlations (Miller & Chapman, 2001). We therefore report analyses with and without FSIQ as a covariate.

For information purposes only, supplementary Table S2 describes the neuropsychological performance of the final sample of patients versus controls. Available pre-treatment results suggest that, as expected, patients may have experienced difficulties in working memory and concentration as previously reported in pre-treatment cancer patients(Cimprich et al., 2005). Also expected was the absence of group differences on memory measures prior to treatment. Notably, the forgetting scores that we report below were computed to minimise the influence of executive difficulties at the time of encoding by computing them relative to baseline scores.

### Delayed forgetting (a measure of slow consolidation)

All participants forgot over one day. Figure 3 depicts asignificant interaction between Session and Group (F_1,18_ =7, p=.02), the only significant effect obtained in this analysis. This interaction remained significant when controlling for FSIQ (F_1,17_=7.20, p=.02). Patients forgot significantly faster than controls after treatment (t_18_=2.64, p.032; g=1.13, 95% CI= .22 to 2.04). Patients also forgot more after treatment than before treatment (t_9_=2.12, p=.031; g=.75, 95% CI = -.11 to 1.62). Individual participant data, depicted in Figure 3, shows that the patients’ accelerated forgetting after treatment was not a result of outlier results. The same results were obtained in the extended sample of N=13 patients and their controls. This finding confirms our key hypothesis that the treatment delivered between the learning session and the delayed retrieval session would result in accelerated forgetting.

**Figure 3.**
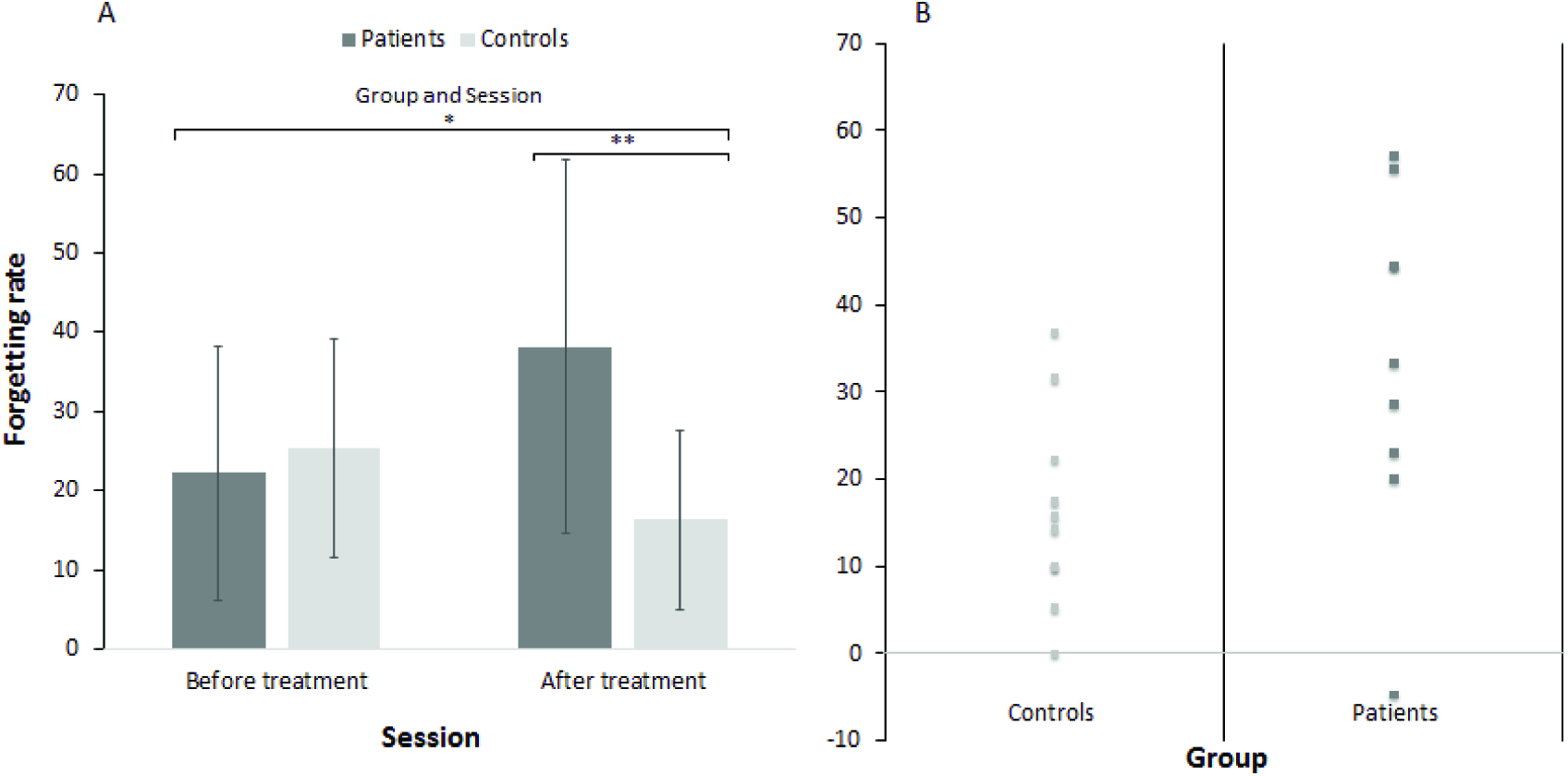
Forgetting rates in patients and controls before and after treatment. A. Error bars represent the standard deviation. B. Scatter plot depicting the percentage of words forgotten by controls and patients (light/dark grey) on Session 3 (after treatment). Each data point represents the individual forgetting rate of a participant.*p<.05, **p<.01.

### Immediate forgetting (a measure of rapid consolidation)

There was no forgetting across the 2-minute distractor task that separated the second and the third free recall tests in both the N=10 or N=13 samples; performance on the later test was numerically higher than performance on the earlier test, possibly due to retrieval practice (Nunes & Karpicke, 2015). No other effects were significant.

### Learning

There was a significant Group effect (F_1,18_=5.10, p=.03), indicative of learning difficulties in patients, which was maintained when including FSIQ as a covariate (F_1,17_=4.30, p=.05). The same results were obtained in the extended sample of N=13, but after controlling for FSIQ the Group effect was only marginally significant, p=0.054. Note that forgetting scores could not be affected by these group differences in learning (see Methods). We have conducted additional analysis of the first and second learning trials in Session 1, and the first and second learning trials in the Sessions before and after treatment (Sessions 2 and 3). These analyses continued to show a main effect of Group which did not interact with any of the other factors. Hence, while the performance of patients, across learning trials, was generally lower than that of controls in all three sessions, we do not have evidence that patients failed to benefit to the same degree as controls from additional learning and retrieval experiences.

**Figure 4.**
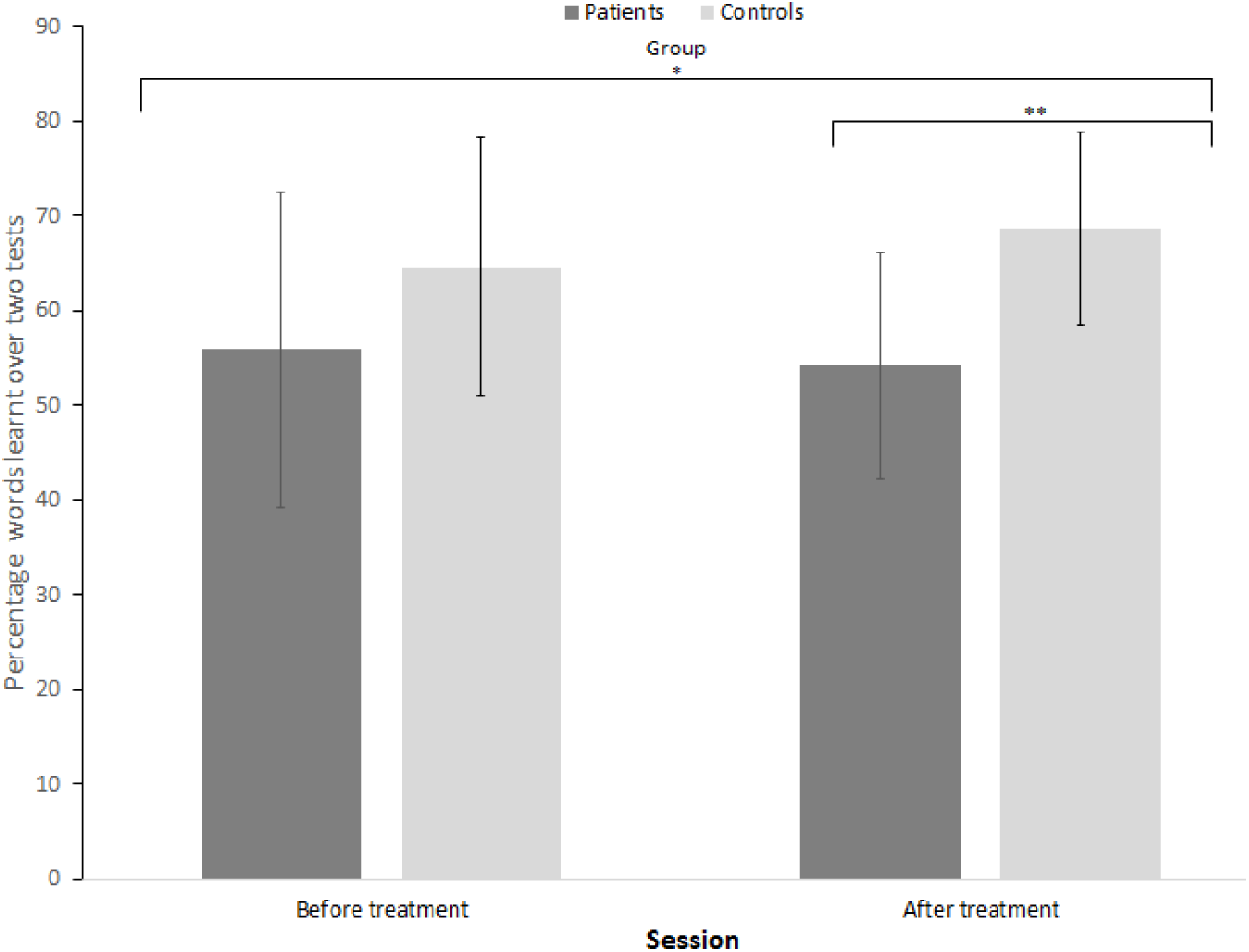
Learning rates in patients and controls before and after treatment. A. Differences between delayed free and cued recall. B. Differences in the average number of intrusions over all recall trials, between patients and controls, before and after treatment. Error bars represent the standard deviation.

### Delayed retrieval

None of the groups performed at ceiling, although all participants benefited from cues, suggesting that the cues were appropriate to the task but none of the effects were significant. The same results were obtained in the extended sample of N=13, although the interaction between Group and Session was marginally significant before controlling for FSIQ, p=0.06.

**Figure 5.**
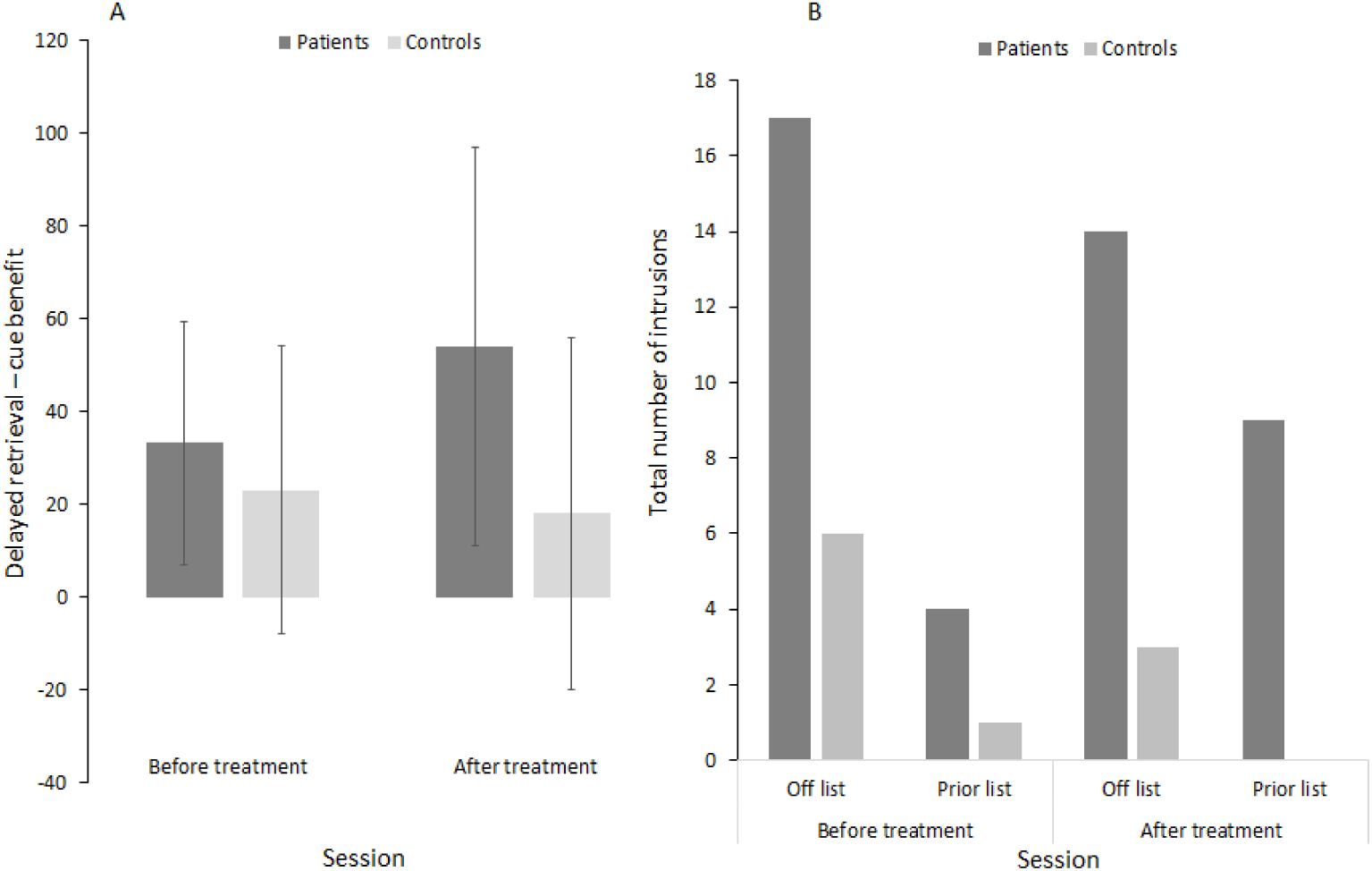
Retrieval performance in patients and controls before and after treatment. Spatial variation of the mean effects of migration (A) and persistence (B) mechanisms on the current climatic debt observed in plant communities throughout the French forests, and balance between both effects (C). (D) Synthesis of species’ migration and persistence effects in each biogeographic region encountered in France. Mean effects (and confidence intervals in panel D) are computed from 1000 bootstrapped models. More negative and positive values indicate higher mitigating and amplifying effects on the climatic debt, respectively.

### Intrusions (a measure of Retrieval)

Patients produced more intrusions than controls, a difference that was significant after treatment (U=16.5, p<.01), but not before treatment (U=35.5, p=.28). The same results were obtained in the extended sample of N=13. This may suggest a retrieval deficit, which was not captured by our retrieval score.

## Discussion

This is the first investigation of acute memory impairments in nCNS cancer patients. We report the results of a small study investigating learning, forgetting rates, and retrieval in patients before and after their first dose of chemotherapy treatment. Recruiting to this cognitive study within weeks of diagnosis presented formidable challenges, so that over a period of three years we were only able to recruit a modest sample of 10 patients. Despite the small sample, the study did reveal three significant differences between patients and controls.

Our key finding was a treatment-related modulation of forgetting. After treatment, patients forgot more over a 24-hour period compared to controls, as measured by free recall. We also observed a significant increase in the number of intrusions patients produced after treatment. Finally, patients performed less well than controls on immediate free recall tests both before and after treatment. We discuss these results below in light of the recruitment challenges and reflect on how these could be overcome in future studies.

Across both pre- and post-treatment testing sessions, patients’ ability to learn and immediately recall what they have learned was poorer compared to that of controls. Previous work has established that cognitive performance in cancer patients prior to their treatment is decreased compared to controls (Ahles et al., 2008), with documented poorer attention, executive functioning, or working memory (Cimprich et al., 2005; Menning et al., 2015). These deficits could be caused by cancer-related frontal dysfunction (Rabbitt, Lowe, & Shilling, 2001) or contextual, transient concentration problems due to anxiety (Hermelink et al., 2015). Anxiety and post-traumatic stress, in particular, have been documented in post-treatment cancer patients (Hermelink et al., 2015; Stark et al., 2002), and vary in prevalence and severity along the disease trajectory (Traeger, Greer, Fernandez-Robles, Temel, & Pirl, 2012). Some subclinical, contextual anxiety isexpected before treatment (Traeger et al., 2012), but, in our sample of patients, anxiety was not particularly increased, so it is a less likely cause for executive deficits. In summary, the learning difficulties patients demonstrated were independent of treatment and are likely related to deficits in executive functions, which have been previously documented in post-treatment patients. Importantly, our key measure of delayed forgetting was independent of initial learning ability, because forgetting rates were computed relative to immediate memory performance, thus controlling for baseline performance differences.

Our main finding was that, in agreement with our hypothesis, delayed forgetting over the course of 24 hours was increased in patients after their first treatment compared to controls and their own pre-treatment performance. The fact that we have demonstrated significantly accelerated forgetting in a relatively under-powered study suggests that the true effect size in the population could be large.This is an important finding because delayed forgetting is thought to be a marker of slow long-term memory consolidation (Hardt et al., 2013). We could not assess changes in rapid consolidation because our participants did not forget across a short 2-minute delay. Our finding suggests that cancer chemotherapy may have acute effects on slow consolidation in humans. This is the first time that cancer chemotherapy has been shown to have acute cognitive effects and that a drug treatment has been shown to accelerate human forgetting. If our findings are corroborated in future studies they could open up a new research avenue that will throw light on the nature of human long-term memory consolidation.

We used a demanding memory test – free recall, because it provides the best markerof accelerated forgetting in patients with hippocampal damage (Isaac & Mayes, 1999a, 1999b). Yet the taxing nature of the task could lead to higher forgetting rates in the patient group due to potential retrieval difficulties, namely difficulties in the executive control of the retrieval process, rather than impaired consolidation that gives rise to weaker memory traces. To distinguish between these two possibilities, the task included a measure of the retrieval process (the degree to which cues facilitated retrieval of the studied material). This measure did not differ between patients and controls, supporting the interpretation that accelerated forgetting was caused by impaired slow consolidation rather than retrieval difficulties; however, given the small sampleas well as the increased number of patient intrusions, null effects should be interpreted with caution. The second retrieval measure we incorporated (number of intrusions), demonstrated a treatment-dependent increase in patients, indicative of potential executive control problems after treatment. If patients suffered from impaired executive control after treatment they may have found it more difficult to access memory traces, even if they were preserved. Differentiating between the two types of deficits, if at all possible, would require a different experimental design with hypotheses that would build up on our findings.

Taken together, increased retrieval difficulties after treatment, possibly related to impaired executive functions, cannot be ruled out as an explanation ofthe observed accelerated forgetting after treatment, indicating the need for future work on this topic. In fact, recent work suggests that poor consolidation and retrieval difficulties are perhaps more closely intertwined than previously thought. Studies in animal models suggests that protein synthesis inhibition, which for many years was thought to cause amnesia by impairing slow consolidation, may not damage the engram cells themselves, but makes it more difficult for organisms to access those memories during memory tests (Ryan, Roy, Pignatelli, Arons, & Tonegawa, 2015; Tonegawa, Pignatelli, Roy, & Ryan, 2015).

In conclusion, our findings indicate that patients forget faster than controls after their first treatment either because of treatment-dependent impairment of slow consolidation or because of retrieval difficulties associated with poorer executive functioning. Future studies should take special care to differentiate between disruptions to synaptic consolidation in the medial temporal lobe, and frontally-mediated retrieval problems. This could be explored in imaging studies by adapting our task to compare changes in free and cued recall in immediate versus delayed tests and by measuring attention-related imaging markers continuously throughout the task. Future animal work could build on our findings to decide whether both frontal and medial temporal lobe regions are affected following the first treatment through a disruption of de novo protein synthesis, necessary in slower consolidation processes.

Other factors could be considered in interpreting the accelerated forgetting and the larger number of intrusions in patients following treatment. First, these findings could be due to increased emotional distress and fatigue after treatment (Cimprich et al., 2005). Our data do not allow us to completely rule out this interpretation because many of our patients failed to return the self-assessment questionnaires aimed at accounting for these effects. That said, there is no a-priori reason to believe that patients were *more* distressed during the post-treatment Session 3 than during the pre-treatment Session 2. Future studies should employ frequent measures of psychological distress. Second, these findings could be related to concomitantly administered medication, such as corticosteroids, administered for antiemetic prophylaxis and known to have effects on medial temporal lobe structures as well as consolidation (Brown, 2009). This possibility is unlikely because post-encoding administration of cortisol is known to *attenuate* rather than accelerate forgetting (McGaugh, 2002). Finally, these findings could be due to the change of physiological context between the Sessions. It is possible that memory for materials studied in Session 2 has been associated with the treatment that was delivered soon afterwards, making it more difficult for patients than controls to access Session 2 materials during Session 3 (Gisquet-Verrier et al., 2015; McCullough & Yonelinas, 2013). This possibility is less likely given that the physiological effects of the drugs would have continued to affect patients during Session 3. Putative effects of context shifts could be evaluated in future studies by administering a fourth Session and checking whether forgetting is accelerated also between Session 3 and Session 4. Larger studies, with a more homogenous treatment regimen, would also help assess context effects by utilizing designs that are informed by the time-course of drug action.

Significant restrictions were imposed from the outset by recruiting patients at an emotionally stressful time, after diagnosis and before treatment onset. They were also posed by the general difficulties in recruiting patients to psycho-social oncology studies in the UK (Ashley et al., 2012), especially for cognitive studies (Shilling, Jenkins, Fallowfield, &Howell, 2003). Recruitment difficulties meant that despite our best efforts over the course of three years, we only achieved a final sample of 10 patients. Therefore, although we attempted to evaluate patients’ neuropsychological status and adhere to the recommendations of the ICCTF (Wefel et al., 2011), not all of our patients completed the battery, and our interpretation of the memory deficits we observed relies on the pattern of patients’ performance on our central memory task, which had its own in-built controls. Recruitment challenges could be overcome in future research by investing more in raising awareness about the scientific evidence for cancer and treatment-dependent cognitive impairments in survivors. For us, recruitment difficulties were only partly overcome by our simplified and brief testing procedure and the use of a flexible computerised task. Future studies could look at further simplifying the delivery of tests by using pencil-and-paper instruments or online assessments, depending on the demographic of the cancer group. Although our sample size does not detract, but rather underscores, the magnitude of the significant effects we obtained, it does mean that we cannot interpret null effects. Clearly, it is critically important to replicate our findings in more highly powered studies.

Despite their limitations, our results are unique in highlighting treatment-related cognitive deficitsin working age cancer patients relative to controls. We hope that our results will encourage others to pursue large sample investigations of memory processes that will pinpoint the biological mechanisms underlying acute and long-term structural brain treatment-related change in nCNS cancer patients, and encourage the development of strategies to mitigate these effects on survivors’ lives.

## Acknowledgements

This work was supported by the Medical Research Council (G1000399). The authors thank all participants and NHS Trusts who contributed to this study. We also thank M Cohn, G Winocur and L Fuentemilla, as well as the reviewers for their valuable comments on our manuscript.

## Supplementary material

**Table S1.** Demographic characteristics of included and excluded patients.

|  | Included patients<br>N, %/M (SD) | Excluded patients<br>N, M (SD) | P |
| --- | --- | --- | --- |
| Age | 10, 28.1 (12.44) | 20, 34.1 (13.6) | .25 |
| Sex | 10, 70% Female | 20, 75% female | NA |

**Table S2.**
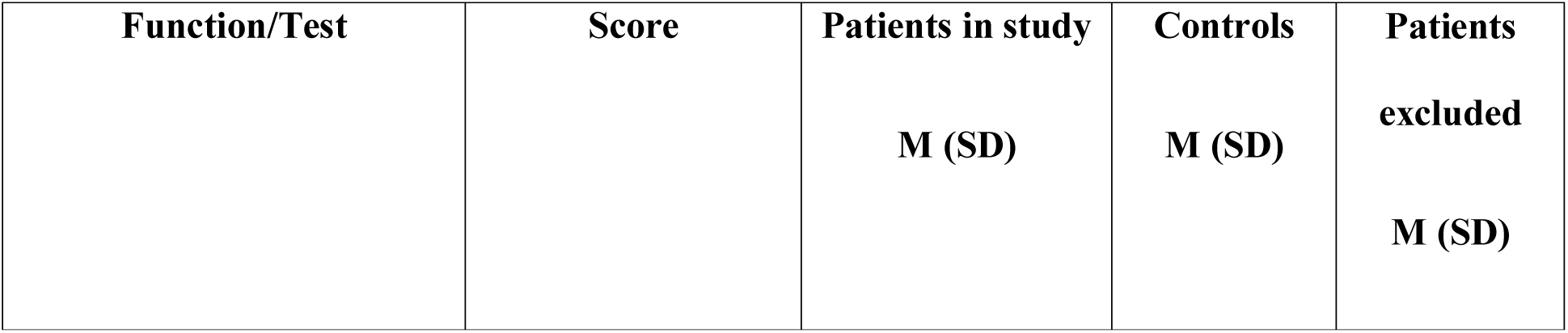
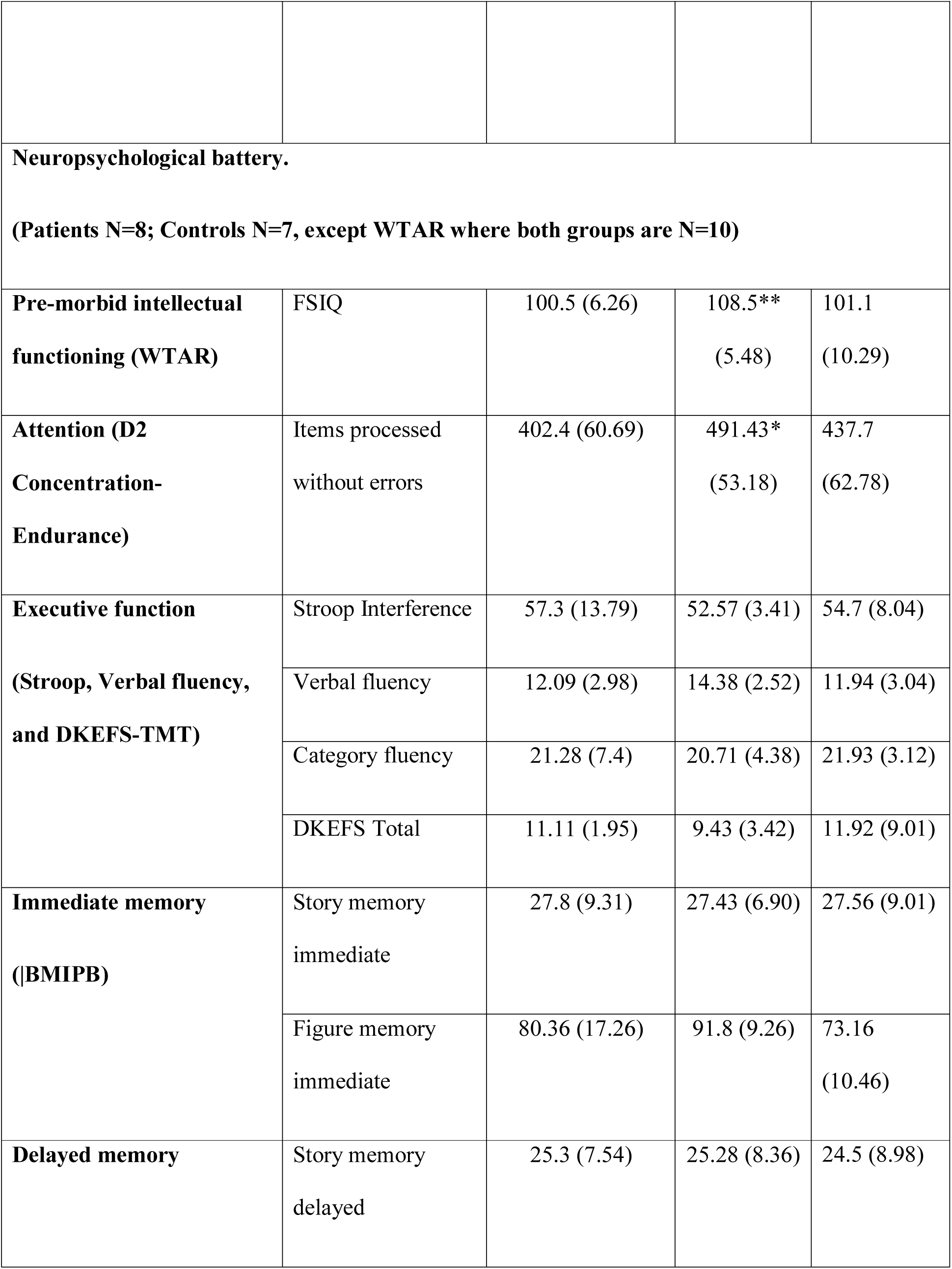
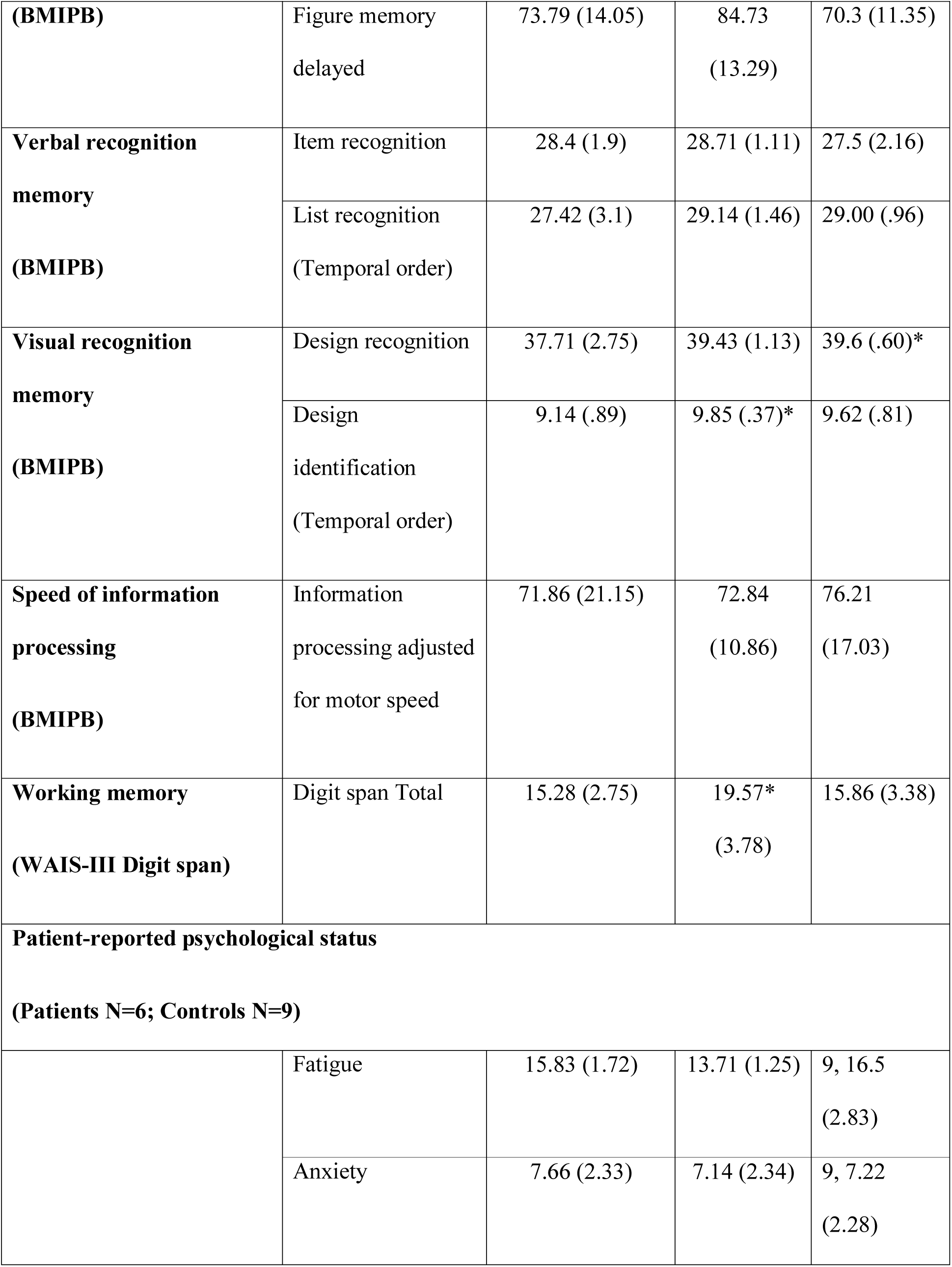
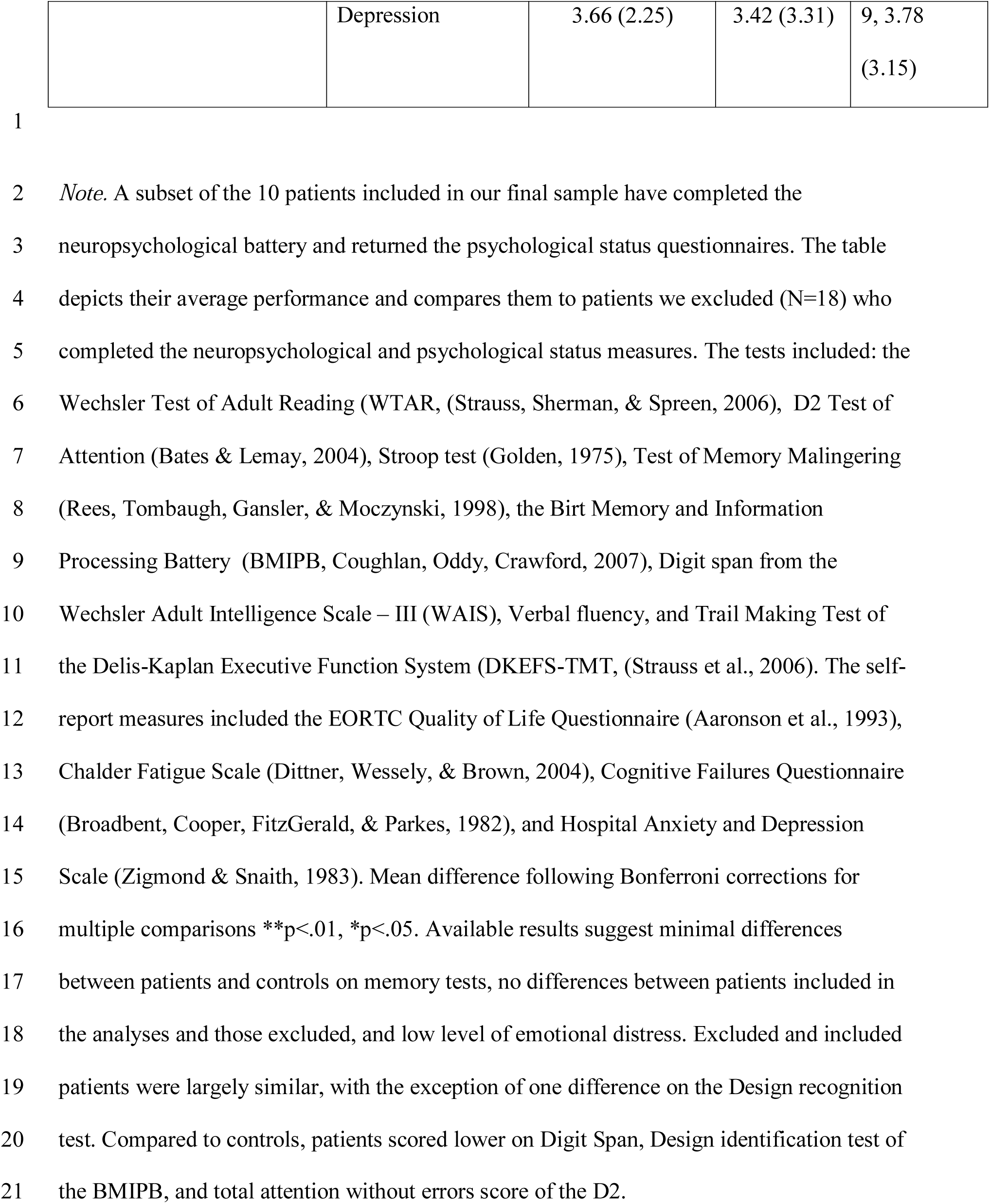
Neuropsychological performance and patient-reported psychological status in patients and controls.

**Table S3.** Characteristics of words used in memory task.

| List | Word | Familiarity | Kucera-Francis<br>Frequency | Summary |
| --- | --- | --- | --- | --- |
| **Animals &**<br>**Vehicles** | **Camel** | 421 | 1 | Familiarity<br>M= 340.95<br>SD=231.02<br>Frequency<br>M=20.21<br>SD=35.6 |
|  | **Helicopter** | 0 | 1 |  |
|  | **Motorcycle** | 0 | 0 |  |
|  | **Elephant** | 459 | 7 |  |
|  | **Bicycle** | 0 | 5 |  |
|  | **Butterfly** | 481 | 2 |  |
|  | **Donkey** | 0 | 1 |  |
|  | **Lion** | 511 | 17 |  |
|  | **Train** | 548 | 82 |  |
|  | **Plane** | 558 | 114 |  |
|  | **Snail** | 489 | 1 |  |
|  | **Rabbit** | 523 | 11 |  |
|  | **Sheep** | 507 | 23 |  |
|  | **Bear** | 526 | 57 |  |
|  | **Ostrich** | 358 | 0 |  |
|  | **Horse** | 560 | 117 |  |
|  | **Sledge** | 0 | 0 |  |
|  | **Squirrel** | 511 | 1 |  |
|  | **Giraffe** | 381 | 0 |  |
|  | **Swan** | 0 | 3 |  |
|  | **Fish** | 548 | 35 |  |
|  | **Kangaroo** | 0 | 0 |  |
|  | **Goat** | 469 | 6 |  |
|  | **Zebra** | 333 | 1 |  |
| **Fruits &**<br>**Clothes** | **Dress** | 588 | 67 | Familiarity<br>M=439.79<br>SD=232.00<br>Frequency<br>M=12.58<br>SD=15.99 |
|  | **Watermelon** | 0 | 0 |  |
|  | **Mitten** | 0 | 0 |  |
|  | **Jumper** | 0 | 1 |  |
|  | **Strawberry** | 539 | 0 |  |
|  | **Pear** | 567 | 6 |  |
|  | **Apple** | 598 | 9 |  |
|  | **Trousers** | 0 | 7 |  |
|  | **Pineapple** | 489 | 9 |  |
|  | **Orange** | 567 | 23 |  |
|  | **Lemon** | 518 | 18 |  |
|  | **Grapes** | 0 | 0 |  |
|  | **Cherry** | 514 | 6 |  |
|  | **Glove** | 575 | 9 |  |
|  | **Necklace** | 536 | 3 |  |
|  | **Banana** | 576 | 4 |  |
|  | **Shoe** | 569 | 14 |  |
|  | **Ring** | 589 | 47 |  |
|  | **Belt** | 550 | 29 |  |
|  | **Boot** | 566 | 13 |  |
|  | **Skirt** | 551 | 21 |  |
|  | **Sock** | 578 | 4 |  |
|  | **Umbrella** | 511 | 8 |  |
|  | **Tomato** | 574 | 4 |  |
| **Vegetables**<br>**& Kitchen**<br>**objects** | **Asparagus** | 534 | 1 | Familiarity<br>M=493.83<br>SD=193.01<br>Frequency<br>M=28.63<br>SD=46.34 |
|  | **Carrot** | 539 | 1 |  |
|  | **Spoon** | 612 | 6 |  |
|  | **Fridge** | 0 | 0 |  |
|  | **Corn** | 548 | 34 |  |
|  | **Glass** | 611 | 99 |  |
|  | **Table** | 599 | 198 |  |
|  | **Iron** | 555 | 43 |  |
|  | **Scissors** | 559 | 1 |  |
|  | **Toaster** | 520 | 0 |  |
|  | **Chair** | 617 | 66 |  |
|  | **Oven** | 577 | 7 |  |
|  | **Mushroom** | 0 | 2 |  |
|  | **Onion** | 550 | 15 |  |
|  | **Pepper** | 554 | 13 |  |
|  | **Potato** | 612 | 15 |  |
|  | **Kettle** | 551 | 3 |  |
|  | **Knife** | 573 | 76 |  |
|  | **Ladder** | 507 | 19 |  |
|  | **Bottle** | 591 | 76 |  |
|  | **Pumpkin** | 0 | 2 |  |
|  | **Broom** | 547 | 2 |  |
|  | **Stool** | 531 | 8 |  |
|  | **Lettuce** | 565 | 0 |  |
| **Four-**<br>**legged**<br>**animals**<br>**and**<br>**Musical**<br>**instruments** | **Alligator** | 442 | 4 | Familiarity<br>M=426.83<br>SD=170.33<br>Frequency<br>M=8.33<br>SD=9.95 |
|  | **Deer** | 509 | 13 |  |
|  | **Frog** | 507 | 1 |  |
|  | **Horn** | 498 | 31 |  |
|  | **Leopard** | 431 | 0 |  |
|  | **Violin** | 468 | 11 |  |
|  | **Monkey** | 531 | 9 |  |
|  | **Accordion** | 394 | 1 |  |
|  | **Turtle** | 509 | 8 |  |
|  | **Racoon** | 0 | 0 |  |
|  | **Trumpet** | 490 | 7 |  |
|  | **Bull** | 0 | 14 |  |
|  | **Ferret** | 0 | 1 |  |
|  | **Guitar** | 550 | 19 |  |
|  | **Rhinoceros** | 400 | 3 |  |
|  | **Flute** | 496 | 1 |  |
|  | **Gorilla** | 554 | 0 |  |
|  | **Seal** | 482 | 17 |  |
|  | **Harp** | 430 | 1 |  |
|  | **Piano** | 545 | 38 |  |
|  | **Tiger** | 513 | 7 |  |
|  | **Beaver** | 470 | 3 |  |
|  | **Skunk** | 519 | 0 |  |
|  | **Drum** | 506 | 11 |  |
| **Birds and**<br>**Toys** | **Chicken** | 544 | 37 | Familiarity<br>M=411.66<br>SD=198.12<br>Frequency<br>M=35.54<br>SD=119.56 |
|  | **Sailboat** | 0 | 0 |  |
|  | **House** | 600 | 591 |  |
|  | **Penguin** | 360 | 0 |  |
|  | **Rooster** | 385 | 3 |  |
|  | **Swing** | 0 | 24 |  |
|  | **Sparrow** | 523 | 0 |  |
|  | **Parrot** | 0 | 1 |  |
|  | **Crow** | 490 | 2 |  |
|  | **Wagon** | 443 | 55 |  |
|  | **Football** | 565 | 36 |  |
|  | **Eagle** | 465 | 5 |  |
|  | **Skate** | 534 | 1 |  |
|  | **Duck** | 529 | 9 |  |
|  | **Dove** | 415 | 4 |  |
|  | **Whistle** | 505 | 4 |  |
|  | **Balloon** | 520 | 10 |  |
|  | **Cannon** | 498 | 7 |  |
|  | **Clown** | 511 | 3 |  |
|  | **Stork** | 393 | 0 |  |
|  | **Kite** | 481 | 1 |  |
|  | **Truck** | 620 | 57 |  |
|  | **Pigeon** | 499 | 3 |  |
|  | **Snowman** | 0 | 0 |  |

*Note.* The unequal distribution of the words in the lists was determined by the number of concepts in the database, which complied with our length, familiarity, and frequency constraints. There were no significant differences between the familiarity and frequency of words in different lists, List Familiarity $F_{4,115}=1.71$ ( $p=.15$ ); List Frequency $F_{4,115}=.83$ ( $p=.51$ )

**Table S4.** Percentage recall on immediate and delayed free recall (FR) and delayed cued recall (CR) tests in each Session and Group before controlling for FSIQ.

| **Session** | **Group** | **FR**<br>**M (SD)** | **FR**<br>**M (SD)** | **FR**<br>**M (SD)** | **FR**<br>**Delayed**<br>**M (SD)** | **CR**<br>**Delayed**<br>**M (SD)** |
| --- | --- | --- | --- | --- | --- | --- |
| **Session 1** | Patient | 46.66<br>(13.58) | 61.66<br>(18.08) | 66.24<br>(14.08) | 52.07<br>(18.34) | 67.08<br>(18.05) |
|  | Control | 61.59 | 77.49 (9.25) | 87.07 (9.90) | 64.57 | 76.66 |
|  |  | (12.97) |  |  | (11.82) | (11.32) |
| **Session 2** | Patient | 47.49 | 64.16 | 68.33 | 42.91 | 62.08 |
|  |  | (18.65) | (14.72) | (18.13) | (21.16) | (26.38) |
|  | Control | 57.49 | 71.66 | 79.99 | 67.08 | 74.99 |
|  |  | (13.29) | (13.99) | (15.69) | (17.72) | (15.08) |
| **Session 3** | Patient | 48.33 | 60 (13.91) | 61.66 |  |  |
|  |  | (10.05) |  | (16.53) |  |  |
|  | Control | 59.74 | 77.49 (8.82) | 87.07 |  |  |
|  |  | (11.62) |  | (11.52) |  |  |

Abbreviations: M=mean; SD=standard deviation

**Figure S1.**
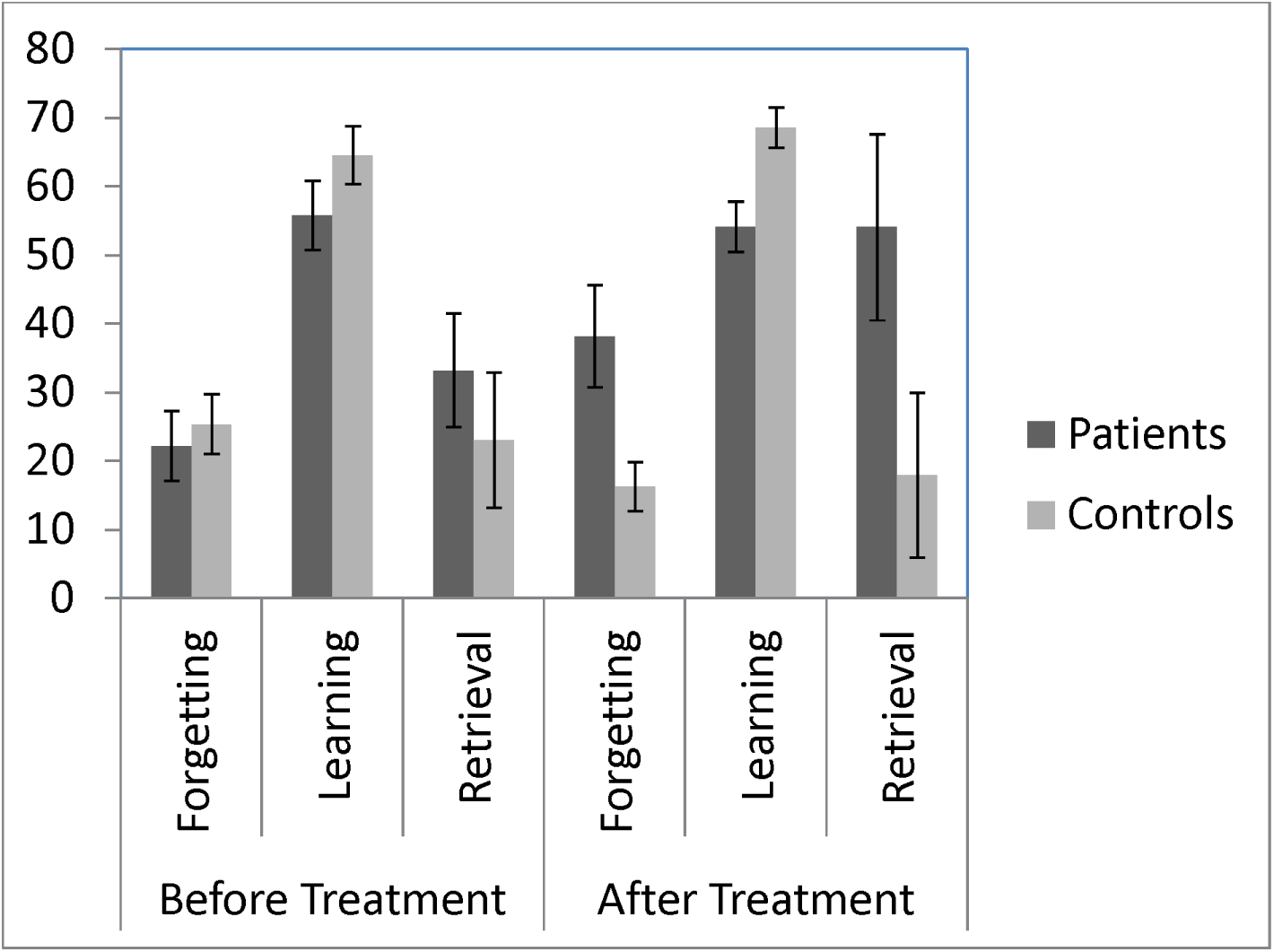
A comparison of forgetting, learning and retrieval scores of the 10 patients included in the manuscript (Top) and an extended sample of 13 patients, including 3 who were tested without a computer in Session 3 (Bottom). The error bars represent standard error.

## Appendix: Memory task instructions

### Session 1

Participants were given the following instructions before they studied the first list of words (List 1):

> “On the screen you are going to see a list of words that I’d like you to remember. The list is quite long, so don’t try to remember all the words from the beginning. That is why we are going to go through the same list several times, for you to be able to remember more words each time. Whenever you see a word on the screen please tell me a sentence containing that word. The sentence can be simple (such as “Cats have fur”, if the word on the screen is “Cat”), but please make sure you use a different sentence for each word. After you see the entire list I will ask you to tell me what you remember from it. The list of words for today will be made out of categories A and B”.

List 1 was presented on the screen, following which participants were asked to Free recall the words they had learned in any order, whilst being reminded of the categories to which they belonged:

> “Now tell me all the words you can remember, in any order”.

If participants paused for more than 20 seconds they were asked:

> “Is that all you can remember?”

Upon confirming, FR11 was concluded, and the following instruction was given:

> “Now we are going to go through the same list of words again just as we did before. Again, tell me a sentence with each word. You can use the same sentence as before”.

The second on-screen presentation of List 1 commenced, which was followed by the second Free recall test (FR12). Using the same instruction, after the test, they were shown the same list for the third time, which was followed by the distracter task. The instructions for the distracter task stated:

> “Before you tell me what you remember, you will play a short game. On the screen, you are going to see two pictures and I want you to tell me how many differences you see between them. You will have 2 minutes to tell me as many differences as you can”.

All the differences participants spotted during the task were recorded by the experimenter. At the end of the 2-minute delay participants were given the same free recall instructions as before:

> “Now tell me all the words you can remember, in any order”.

If participants paused for more than 20 seconds they were asked:

> “Is that all you can remember?”

If they confirmed, the Session was concluded.

### Session 2

Participants were first administered a surprise free delayed test (FR14) for the list they studied during Session 1. The instructions for that test were:

> “Could you tell me what words you remember off the list of Categories A and B you learned yesterday?

They were allowed to Free recall at their own pace, and the test was concluded if they could not remember any more words for 20 seconds, as in the previous tests.

They were then administered a surprise cued recall test of the same list (CR15). The instructions for that test were:

> “Now we are going to do something a bit different. On the screen you are going to see the first two letters of each of the words you learned in this list. They may help you remember a few more words. Don’t think about it for too long – if the word comes immediately to mind, tell me what it is. If it doesn’t, just say Pass.”

A second List of words was studied in a process identical to the one in Session 1, whilst displaying the words on the screen:

> “Now we are going to go through another list of words just as we did yesterday – three consecutive times and with sentences. This time the list of words that you will be learning is made out of Categories A and B”.

The remainder of Session 2 had the same instruction as Session 1, while Session 3 had the same instructions as Session 2.

## References

Ahles, T. A, & Saykin, A. J. (2007). Candidate mechanisms for chemotherapy-induced cognitive changes. Nature Reviews. Cancer, 7(3), 192–201. http://doi.org/10.1038/nrc2073

Ahles, T. A, Saykin, A. J., McDonald, B. C., Furstenberg, C. T., Cole, B. F., Hanscom, B. S., … Kaufman, P. A. (2008). Cognitive function in breast cancer patients prior to adjuvant treatment. Breast Cancer Research and Treatment, 110(1), 143–52. http://doi.org/10.1007/s10549-007-9686-5

Alber, J., Della Sala, S., & Dewar, M. (2014). Minimizing interference with early consolidation boosts 7-day retention in amnesic patients. Neuropsychology, 28(5), 667–675. http://doi.org/10.1037/neu0000091

Arruda-Carvalho, M., Sakaguchi, M., Akers, K. G., Josselyn, S. A., & Frankland, P. W. (2011). Posttraining ablation of adult-generated neurons degrades previously acquired memories. The Journal of Neuroscience□: The Official Journal of the Society for Neuroscience, 31(42), 15113–27. http://doi.org/10.1523/JNEUROSCI.3432-11.2011

Ashley, L., Jones, H., Velikova, G., & Wright, P. (2012). Cancer patients’ and clinicians’ opinions on the best time in secondary care to approach patients for recruitment to longitudinal questionnaire-based research. Supportive Care in Cancer, 20(12), 3365–3372. http://doi.org/10.1007/s00520-012-1518-4

Averell, L., & Heathcote, A. (2011). The form of the forgetting curve and the fate of memories. Journal of Mathematical Psychology, 55(1), 25–35. http://doi.org/10.1016/j.jmp.2010.08.009

Baddeley, A., & Wilson, B. Frontal amnesia and the dysexecutive syndrome., 7Brain and cognition 212–230 (1988). http://doi.org/10.1016/0278-2626(88)90031-0

Borenstein, M., Hedges, L. V., Higgins, J. P. T., & Rothstein, H. R. (2009). Introduction to Meta-Analysis. Chichester, UK: John Wiley & Sons, Ltd. http://doi.org/10.1002/9780470743386

Brearley, S. G., Stamataki, Z., Addington-Hall, J., Foster, C., Hodges, L., Jarrett, N., … Amir, Z. (2011). The physical and practical problems experienced by cancer survivors: A rapid review and synthesis of the literature. European Journal of Oncology Nursing, 15(3), 204–212. http://doi.org/10.1016/j.ejon.2011.02.005

Brown, E. S. (2009). Effects of glucocorticoids on mood, memory, and the hippocampus. Treatment and preventive therapy. Annals of the New York Academy of Sciences, 1179, 41–55. http://doi.org/10.1111/j.1749-6632.2009.04981.x

Carlesimo, G. A., Cherubini, A., Caltagirone, C., & Spalletta, G. (2010). Hippocampal mean diffusivity and memory in healthy elderly individuals: A cross-sectional study. Neurology, 74(3), 194–200. http://doi.org/10.1212/WNL.0b013e3181cb3e39

Cimprich, B., So, H., Ronis, D. L., & Trask, C. (2005). Pre-treatment factors related to cognitive functioning in women newly diagnosed with breast cancer. Psycho-Oncology, 14(1), 70–78. http://doi.org/10.1002/pon.821

Coussens, L. M., & Werb, Z. (2002). Inflammation and cancer. Nature, 420(6917), 860–7. http://doi.org/10.1038/nature01322

CRUK. (2014). Cancer survival by age. Retrieved from http://www.cancerresearchuk.org/cancer-info/cancerstats/survival/age/

da Ros, M., Iorio, A. L., Lucchesi, M., Stival, A., de Martino, M., & Sardi, I. (2015). The Use of Anthracyclines for Therapy of CNS Tumors. Anti-Cancer Agents in Medicinal Chemistry, 15(6), 721–727.

de Ruiter, M. B., & Schagen, S. B. (2013). Functional MRI studies in non-CNS cancers. Brain Imaging and Behavior, 7(4), 388–408. http://doi.org/10.1007/s11682-013-9249-9

Deary, I. J., & Johnson, W. (2010). Intelligence and education: causal perceptions drive analytic processes and therefore conclusions. International Journal of Epidemiology, 39(5), 1362–9. http://doi.org/10.1093/ije/dyq072

Deprez, S., Billiet, T., Sunaert, S., & Leemans, A. (2013). Diffusion tensor MRI of chemotherapy-induced cognitive impairment in non-CNS cancer patients: a review. Brain Imaging and Behavior, 7(4), 409–35. http://doi.org/10.1007/s11682-012-9220-1

Dewar, M., Della Sala, S., Beschin, N., & Cowan, N. (2010). Profound Retroactive Interference in Anterograde Amnesia: What interferes? Neuropsychology, 24(3), 357–367. http://doi.org/10.1037/a0018207

Dietrich, J., Prust, M., & Kaiser, J. (2015). Chemotherapy, cognitive impairment and hippocampal toxicity. Neuroscience. http://doi.org/10.1016/j.neuroscience.2015.06.016

Elliott, G., Isaac, C. L., & Muhlert, N. (2014). Measuring forgetting: A critical review of accelerated long-term forgetting studies. Cortex. http://doi.org/10.1016/j.cortex.2014.02.001

Faul, F., Erdfelder, E., Lang, A.-G., & Buchner, A. (2007). G*Power 3: a flexible statistical power analysis program for the social, behavioral, and biomedical sciences. Behavior Research Methods, 39, 175–191. http://doi.org/10.3758/BF03193146

Gershman, S. J., Blei, D. M., & Niv, Y. (2010). Context, learning, and extinction. Psychological Review, 117(1), 197–209. http://doi.org/10.1037/a0017808

Gisquet-Verrier, P., Lynch, J. F., Cutolo, P., Toledano, D., Ulmen, A., Jasnow, A. M., & Riccio, D. C. (2015). Integration of New Information with Active Memory Accounts for Retrograde Amnesia: A Challenge to the Consolidation/Reconsolidation Hypothesis? Journal of Neuroscience, 35(33), 11623–11633. http://doi.org/10.1523/JNEUROSCI.1386-15.2015

Hardt, O., Nader, K., & Nadel, L. (2013). Decay happens: the role of active forgetting in memory. Trends in Cognitive Sciences, 17(3), 111–20. http://doi.org/10.1016/j.tics.2013.01.001

Hermelink, K., Voigt, V., Kaste, J., Neufeld, F., Wuerstlein, R., Bühner, M., … Harbeck, N. (2015). Elucidating pretreatment cognitive impairment in breast cancer patients: the impact of cancer-related post-traumatic stress. Journal of the National Cancer Institute, 107(7), djv099-. http://doi.org/10.1093/jnci/djv099

Hoefeijzers, S., Dewar, M., Della Sala, S., Butler, C., & Zeman, A. (2015). Accelerated long-term forgetting can become apparent within 3-8 hours of wakefulness in patients with transient epileptic amnesia. Neuropsychology, 29(1), 117–25. http://doi.org/10.1037/neu0000114

Isaac, C. L., & Mayes, A. R. (1999a). Rate of forgetting in amnesia: I. Recall and recognition of prose. Journal of Experimental Psychology: Learning, Memory, and Cognition, 25(4), 942–962. http://doi.org/10.1037/0278-7393.25.4.942

Isaac, C. L., & Mayes, A. R. (1999b). Rate of forgetting in amnesia: II. Recall and recognition of word lists at different levels of organization. Journal of Experimental Psychology. Learning, Memory, and Cognition, 25(4), 963–977. http://doi.org/10.1037/0278-7393.25.4.963

Kahana, M., & Adler, M. (2002). Note on the power law of forgetting. … of Pennsylvania, Unpublished Note, (1996), 1–15. Retrieved from https://memory.psych.upenn.edu/files/pubs/KahaAdle02.pdf

Kaiser, J., Bledowski, C., & Dietrich, J. (2014). Neural correlates of chemotherapy-related cognitive impairment. Cortex; a Journal Devoted to the Study of the Nervous System and Behavior, 54, 33–50. http://doi.org/10.1016/j.cortex.2014.01.010

Kandel, E. R., Dudai, Y., & Mayford, M. R. (2014). The molecular and systems biology of memory. Cell. Cell Press.

Kopelman, M. D., Bright, P., Buckman, J., Fradera, A., Yoshimasu, H., Jacobson, C., & Colchester, A. C. F. (2007). Recall and recognition memory in amnesia: Patients with hippocampal, medial temporal, temporal lobe or frontal pathology. Neuropsychologia, 45(6), 1232–1246. http://doi.org/10.1016/j.neuropsychologia.2006.10.005

Lindner, O. C., Phillips, B., McCabe, M. G., Mayes, A., Wearden, A., Varese, F., & Talmi, D. (2014). A meta-analysis of cognitive impairment following adult cancer chemotherapy. Neuropsychology, 28(5), 726–40. http://doi.org/10.1037/neu0000064

Liu, R.-Y., Zhang, Y., Coughlin, B. L., Cleary, L. J., & Byrne, J. H. (2014). Doxorubicin attenuates serotonin-induced long-term synaptic facilitation by phosphorylation of p38 mitogen-activated protein kinase. The Journal of Neuroscience□: The Official Journal of the Society for Neuroscience, 34(40), 13289–300. http://doi.org/10.1523/JNEUROSCI.0538-14.2014

Loftus, G. R., Bolles, R., Hunt, E., Loftus, E., Nelson, T., Nelson, W., … Waggoner, W. (1985). Evaluating Forgetting Curves. Journal of Experimental Psychology: Learning, Memory, and Cognition, 11(2), 397–406. http://doi.org/10.1037//0278-7393.11.2.397

Luu, P., Sill, O. C., Gao, L., Becker, S., Wojtowicz, J. M., & Smith, D. M. (2012). The role of adult hippocampal neurogenesis in reducing interference. Behavioral Neuroscience, 126(3), 381–91. http://doi.org/10.1037/a0028252

MacLeod, C., & Mathews,A. (1988). Anxiety and the allocation of attention to threat. The Quarterly Journal of Experimental Psychology. A, Human Experimental Psychology, 40(4), 653–670. http://doi.org/10.1080/14640748808402292

Mayes, A. R. (1995). Memory and amnesia. Behavioural Brain Research, 66, 29–36. http://doi.org/10.1016/0166-4328(94)00120-5

Mayes, A. R., & Roberts, N. (2001). Theories of episodic memory. Philosophical Transactions of the Royal Society of London. Series B, Biological Sciences, 356(1413), 1395–408. http://doi.org/10.1098/rstb.2001.0941

McCullough, A. M., & Yonelinas, A. P. (2013). Cold-pressor stress after learning enhances familiarity-based recognition memory in men. Neurobiology of Learning and Memory, 106, 11–17. http://doi.org/10.1016/j.nlm.2013.06.011

McGaugh, J. L. (2000). Neuroscience - Memory - a century of consolidation. Science, 287(5451), 248–251. http://doi.org/10.1126/science.287.5451.248

McGaugh, J. L. (2002). Memory consolidation and the amygdala: a systems perspective. Trends in Neurosciences, 25(9), 456. Retrieved from http://www.ncbi.nlm.nih.gov/pubmed/12183206

Menning, S., de Ruiter, M. B., Veltman, D. J., Koppelmans, V., Kirschbaum, C., Boogerd, W., … Schagen, S. B. (2015). Multimodal MRI and cognitive function in patients with breast cancer prior to adjuvant treatment — The role of fatigue. NeuroImage: Clinical, 7, 547–554. http://doi.org/10.1016/j.nicl.2015.02.005

Menon, U., Gentry-Maharaj, A., Ryan, A., Sharma, A., Burnell, M., Hallett, R., … Jacobs, I. (2008). Recruitment to multicentre trials--lessons from UKCTOCS: descriptive study. BMJ (Clinical Research Ed.), 337, a2079. http://doi.org/10.1136/bmj.a2079

Miller, G.A., & Chapman, J. P. (2001). Misunderstanding analysis of covariance. Journal of Abnormal Psychology, 110(1), 40–48. http://doi.org/10.1037/0021-843X.110.1.40

Neisser, U., Boodoo, G., Bouchard, T. J., Boykin, A. W., Ceci, S. J., Loehlin, J. C., & Sternberg, R. J. (1996). Intelligence□: Knowns and Unknowns, 77–101.

Nunes, L. D., & Karpicke, J. D. (2015). Retrieval-based learning: Research at the interface between cognitive science and education. Emerging Trends in the Social and Behavioral Sciences, 1–16.

Pan, W., Stone, K. P., Hsuchou, H., Manda, V. K., Zhang, Y., & Kastin, A. J. (2011). Cytokine signaling modulates blood-brain barrier function. Current Pharmaceutical Design, 17(33), 3729–40.

Pomykala, K. L., de Ruiter, M. B., Deprez, S., McDonald, B. C., & Silverman, D. H. S. (2013). Integrating imaging findings in evaluating the post-chemotherapy brain. Brain Imaging and Behavior, 7(4), 436–52. http://doi.org/10.1007/s11682-013-9239-y

Rabbitt, P., Lowe, C., & Shilling, V. (2001). Frontal tests and models for cognitive ageing. European Journal of Cognitive Psychology. http://doi.org/10.1080/09541440125722

Reese, H. (1997). Counterbalancing and other uses of repeated-measures Latin-square designs: Analyses and interpretations. Journal of Experimental Child Psychology, 158, 137–158.

Richardson, A., Addington-Hall, J., Amir, Z., Foster, C., Stark, D., Armes, J., … Sharpe, M. (2011). Knowledge, ignorance and priorities for research in key areas of cancer survivorship: findings from a scoping review. British Journal of Cancer, 105, 82–94. http://doi.org/10.1038/bjc.2011.425

Rubin, D. C., Wenzel, A. E., Anderson, J., Boneau, A., Cerella, J., Crovitz, H., … Wixted, J. (1996). One Hundred Years of Forgetting□: A Quantitative Description of Retention, 103(4), 734–760.

Ryan, T. J., Roy, D. S., Pignatelli, M., Arons, A., & Tonegawa, S. (2015). Engram cells retain memory under retrograde amnesia. Science, 348(6238), 1007–1013. http://doi.org/10.1126/science.aaa5542

Rzeski, W., Pruskil, S., Macke, A., Felderhoff-Mueser, U., Reiher, A. K., Hoerster, F., … Ikonomidou, C. (2004). Anticancer agents are potent neurotoxins in vitro and in vivo. Annals of Neurology, 56(3), 351–60. http://doi.org/10.1002/ana.20185

Sahay, A., Scobie, K. N., Hill, A. S., O’Carroll, C. M., Kheirbek, M. A., Burghardt, N. S., … Hen, R. (2011). Increasing adult hippocampal neurogenesis is sufficient to improve pattern separation. Nature, 472(7344), 466–470. http://doi.org/10.1038/nature09817

Saykin, A. J., de Ruiter, M. B., McDonald, B. C., Deprez, S., & Silverman, D. H. S. (2013). Neuroimaging biomarkers and cognitive function in non-CNS cancer and its treatment: current status and recommendations for future research. Brain Imaging and Behavior, 7(4), 363–73. http://doi.org/10.1007/s11682-013-9283-7

Seigers, R., & Fardell, J. E. (2011). Neurobiological basis of chemotherapy-induced cognitive impairment: a review of rodent research. Neuroscience and Biobehavioral Reviews, 35(3), 729–41. http://doi.org/10.1016/j.neubiorev.2010.09.006

Shapiro, S. S., Wilk, M. B., & Chen, H. J. (1968). A Comparative Study of Various Tests for Normality. J Am Stat Assoc, 63, 1343–1372. http://doi.org/10.2307/2285889

Shilling, V., Jenkins, V., Fallowfield, L., & Howell, T. (2003). The effects of hormone therapy on cognition in breast cancer. The Journal of Steroid Biochemistry and Molecular Biology, 86(3–5), 405–412. http://doi.org/10.1016/j.jsbmb.2003.07.001

Snodgrass, J. G., & Vanderwart, M. (1980). A standardized set of 260 pictures: norms for name agreement, image agreement, familiarity, and visual complexity. Journal of Experimental Psychology. Human Learning and Memory, 6(2), 174–215. Retrieved from http://www.ncbi.nlm.nih.gov/pubmed/7373248

Stark, D., Kiely, M., Smith, A., Velikova, G., House, A., & Selby, P. (2002). Anxiety disorders in cancer patients: Their nature, associations, and relation to quality of life. Journal of Clinical Oncology, 20(14), 3137–3148. http://doi.org/10.1200/JCO.2002.08.549

Terrando, N., Monaco, C., Ma, D., Foxwell, B. M. J., Feldmann, M., & Maze, M. (2010). Tumor necrosis factor-alpha triggers a cytokine cascade yielding postoperative cognitive decline. Proceedings of the National Academy of Sciences of the United States of America, 107(47), 20518–22. http://doi.org/10.1073/pnas.1014557107

Tonegawa, S., Pignatelli, M., Roy, D. S., & Ryan, T. J. (2015). Memory engram storage and retrieval. Current Opinion in Neurobiology. http://doi.org/10.1016/j.conb.2015.07.009

Traeger, L., Greer, J. A., Fernandez-Robles, C., Temel, J. S., & Pirl, W. F. (2012). Evidence-based treatment of anxiety in patients with cancer. Journal of Clinical Oncology. http://doi.org/10.1200/JCO.2011.39.5632

Vardy, J., Wefel, J. S., Ahles, T., Tannock, I. F., & Schagen, S. B. (2008). Cancer and cancer-therapy related cognitive dysfunction: an international perspective from the Venice cognitive workshop. Annals of Oncology□: Official Journal of the European Society for Medical Oncology / ESMO, 19(4), 623–9. http://doi.org/10.1093/annonc/mdm500

Wefel, J. S., Vardy, J., Ahles, T., & Schagen, S. B. (2011). International Cognition and Cancer Task Force recommendations to harmonise studies of cognitive function in patients with cancer. The Lancet. Oncology, 12(7), 703–8. http://doi.org/10.1016/S1470-2045(10)70294-1

Wixted, J. T. (1990). Analyzing the empirical course of forgetting. Journal of Experimental Psychology: Learning, Memory, and Cognition, 16(5), 927–935. http://doi.org/10.1037/0278-7393.16.5.927

Wixted, J. T. (2004). The psychology and neuroscience of forgetting. Annual Review of Psychology, 55, 235–69. http://doi.org/10.1146/annurev.psych.55.090902.141555

Zhang, C. L., Zou, Y., He, W., Gage, F. H., & Evans, R. M. (2008). A role for adult TLX-positive neural stem cells in learning and behaviour. Nature, 451(7181), 1004–1007. http://doi.org/10.1038/nature06562

